# The phospho-ferrozine assay: A tool to study bacterial redox-active metabolites produced at the plant root

**DOI:** 10.1101/2024.09.04.611340

**Authors:** David Giacalone, Emilly Schutt, Darcy L. McRose

## Abstract

Soil microbial communities are pivotal to plant health and nutrient acquisition. It is becoming increasingly clear that many interactions, both among and between microbes and plants, are governed by small bioactive molecules or “secondary metabolites” that can aid in communication, competition, and nutrient uptake. Yet, secondary metabolite biogeography – who makes what, where, and why— is in its infancy. Further, secondary metabolite biosynthesis genes are often silent or weakly expressed under standard laboratory conditions, making it incredibly difficult to study these small molecules. To begin to address these dual challenges, we focused on Redox-Active metabolites (RAMs), a specific class of small molecules, and took advantage of recent findings that many RAMs aid in acquiring phosphorus and that their production is frequently stimulated by stress for this macronutrient. We developed a screen for RAM-producing bacteria that leverages phosphorus limitation to stimulate metabolite biosynthesis and uses a colorimetric (ferrozine) iron-reduction assay to identify redox activity. We isolated 557 root-associated bacteria from grasses collected at sites across the United States (Santa Rita Experimental Range (AZ), Konza Prairie Biological Station (KS), and Harvard Forest (MA)) and from commercial tomato plants and screened them for RAM production. We identified 128 soil isolates of at least 19 genera across Proteobacteria, Actinobacteria, Firmicutes, and Bacteroidetes that produced RAMs under phosphorus stress. Our work reveals that the production of RAMs under phosphorus stress is common across diverse soil bacteria and provides an approach to screen for these small molecules rapidly.

**Importance:** By secreting secondary metabolites, bacteria at the plant root can defend against diseases and help acquire essential nutrients. However, the genes which synthesize secondary metabolites are typically inactive or are weakly expressed under standard laboratory conditions. This fact makes it difficult to study these small molecules and hinders the discovery of novel small molecules that may play crucial roles in agricultural and biomedical settings. Here, we focus on Redox-Active metabolites (RAMs), a class of secondary metabolites that can help bacteria solubilize phosphorus and are often produced when phosphorus is limited. We developed a screen that rapidly identifies RAM-producing bacteria by utilizing a colorimetric iron-reduction assay in combination with phosphorus limitation to stimulate biosynthesis. The screen reveals that RAM-producing bacteria are far more prevalent in soil than previously appreciated and that this approach can be used to identify RAM producers.

## Introduction

Microbial communities living at the plant root can make important contributions to plant health, and these communities are frequently controlled by exchanges of small molecules or secondary metabolites. These molecules can shape community composition, regulate commensal and pathogenic plant-microbe relationships, and aid in nutrient acquisition (1–7). Although secondary metabolites are increasingly appreciated for their role in microbe-microbe and plant-microbe interactions in soil, the sheer diversity of structures and reactivities is staggering and we lack a fundamental understanding of which metabolites and metabolite producers are most influential in the rhizosphere.

The question of how to identify secondary metabolites that govern microbial communities is a long-standing one across natural and biomedical contexts (8–12). A key part of the problem is that while secondary metabolite biosynthetic gene clusters are ubiquitous in microbial genomes from diverse environments (13–17) most are not produced under standard laboratory conditions (18, 19). The difficulty in activating small molecule biosynthesis is as fascinating as it is frustrating and suggests that a better understanding of the regulation of secondary metabolite biosynthesis might hold the key to understanding the function of these molecules in the wild. Indeed, low doses of antibiotics have emerged as some of the most effective activators of small molecule biosynthesis (18, 20), implying that the production of many secondary metabolites is tuned by growth with other microbes. Nutrient stress can also lead to production of small molecules and in some cases has offered clues as to environmental function. The most notable example of this is the production of siderophores, iron-binding small molecules that are produced under iron limitation and aid bacteria in accessing this nutrient (21).

We recently reported a link between redox-active secondary metabolites and limitation for the macronutrient phosphorus (P) that is analogous to that between siderophores and iron and may help us better understand the function of these molecules in the rhizosphere. Redox-Active Metabolites (RAMs) are a class of small, secreted molecules defined by their ability to perform electron-transfer reactions. One of the best-studied groups of RAMs are molecules called phenazines which are commonly produced by pseudomonads and can take on a variety of roles including antibiotic activity, energy conservation, and nutrient solubilization (22–24). The P-controlled PhoBR two-component system regulates the biosynthesis of phenazines in many species of *Pseudomonas* and is also predicted to regulate the production of secondary metabolites in other bacteria (4, 25). While difficult to understand in infection contexts, this regulatory response may be especially useful in soils where phosphate is often immobilized on the surface of minerals like iron(III) oxides (26). The interaction between phosphate and soil minerals contributes to low P bioavailability across both natural and agricultural systems, often making it difficult for plants to access P even in heavily fertilized soils (27, 28). Owing to their capacity to reduce iron(III) to iron(II), we previously found that RAMs can increase the bioavailability of mineral-associated phosphorus (4). Here, we explore this link between RAMs and P stress as a tool for secondary metabolite discovery and a means to better understand how microbes use these molecules in soils and how this may contribute to plant health.

We reasoned that regulation by P might serve as an expedient way to stimulate RAM production while redox activity could act as a convenient chemical hook that can be easily detected through the interactions of RAMs with iron. If RAMS are poised at an appropriate redox potential, their presence should be detectable by monitoring the reduction of iron(III) to iron(II) via the ferrozine assay, a well-established colorimetric technique for quantifying iron(II) (3, 29, 30) (Fig. 1A). We combined these two ideas to yield the phospho-ferrozine assay which we vetted using synthetic RAMs and cultured pseudomonads. To determine the frequency of P-regulated RAM production in the rhizosphere, we generated a library of 557 root-associated bacterial isolates from sites across the United States. By applying the phospho-ferrozine assay, we found that 28% of isolates across Proteobacteria, Actinobacteria, Firmicutes, and Bacteroidetes likely produce P-regulated RAMs. We focused further genomic and metabolomic investigations on small molecule production in a soil isolate identified as *Pseudomonas kielensis*. Contrary to expectations, this organism does not appear to synthesize a known phenazine but instead makes a different RAM under P stress. Overall, this work highlights the prevalence of P-regulated RAM production in soil bacteria, provides a rapid and efficient approach to screen for this trait, and suggests that RAMs likely play an important role in shaping plant and microbial access to P.

**Figure 1.**
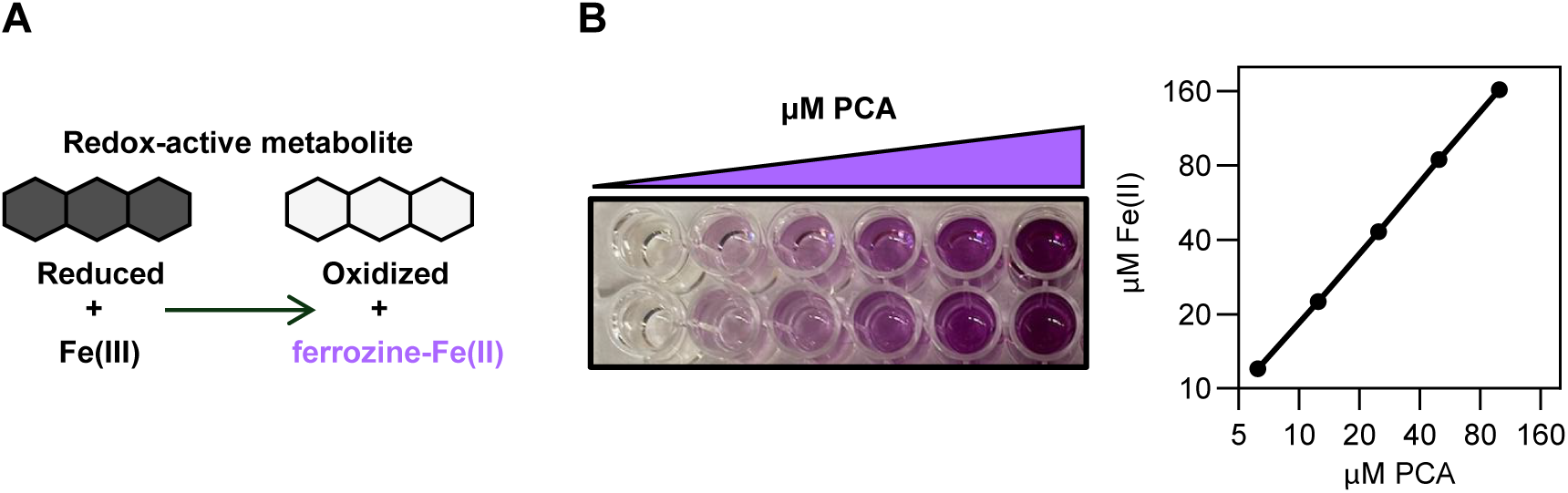
Ferrozine detects iron(III) reduction by redox-active metabolites. **(A)** Schematic showing use of the ferrozine assay to detect redox-active metabolites. RAMs reduce iron(III) to iron(II), ferrozine forms a complex with iron(II) which can be detected via measurement of absorbance at 562 nm. **(B)** An example image and quantification of iron(III) reduction by PCA using ferrozine, showing a ∼2:1 stoichiometry of iron(II):PCA as expected for a two-electron transfer (3). Data shown are the means ± SD of two replicates.

## Results and Discussion

### Development of the phospho-ferrozine assay as a screen for bacteria that produce RAMs

As a proof of concept, we first determined whether the phospho-ferrozine assay could be used to detect the production of RAMs in bacteria known to make these small molecules (Fig. 1A). We chose to focus on *Pseudomonas synxantha* as this bacterium is frequently found in soil and is well studied for the production of phenazines (22, 24, 31), which are synthesized under P-limitation (4, 32–34). We verified that a purified standard of phenazine-1-carboxylic acid (PCA), a RAM made by *P. synxantha* as well as numerous soil-dwelling pseudomonads (35, 36), can facilitate iron reduction and lead to a positive ferrozine signal. Consistent with previous findings, PCA produces a ferrozine signal when mixed with iron(III) (Fig. 1B) (3). Next, we sought to translate our abiotic findings to whole-cell assays. To do so, we aimed to define a P concentration that would stimulate RAM production but maintain robust growth. To ensure that RAM production is specific to P-limitation, we also optimized growth under nitrogen (N) limitation, a condition that is not expected to stimulate RAM production (4). We found that 0.1 mM P and 2 mM N were optimal concentrations to limit bacterial growth given that *P. synxantha* obtained similar optical densities in both conditions (Fig. 2A, purple lines).

**Figure 2.**
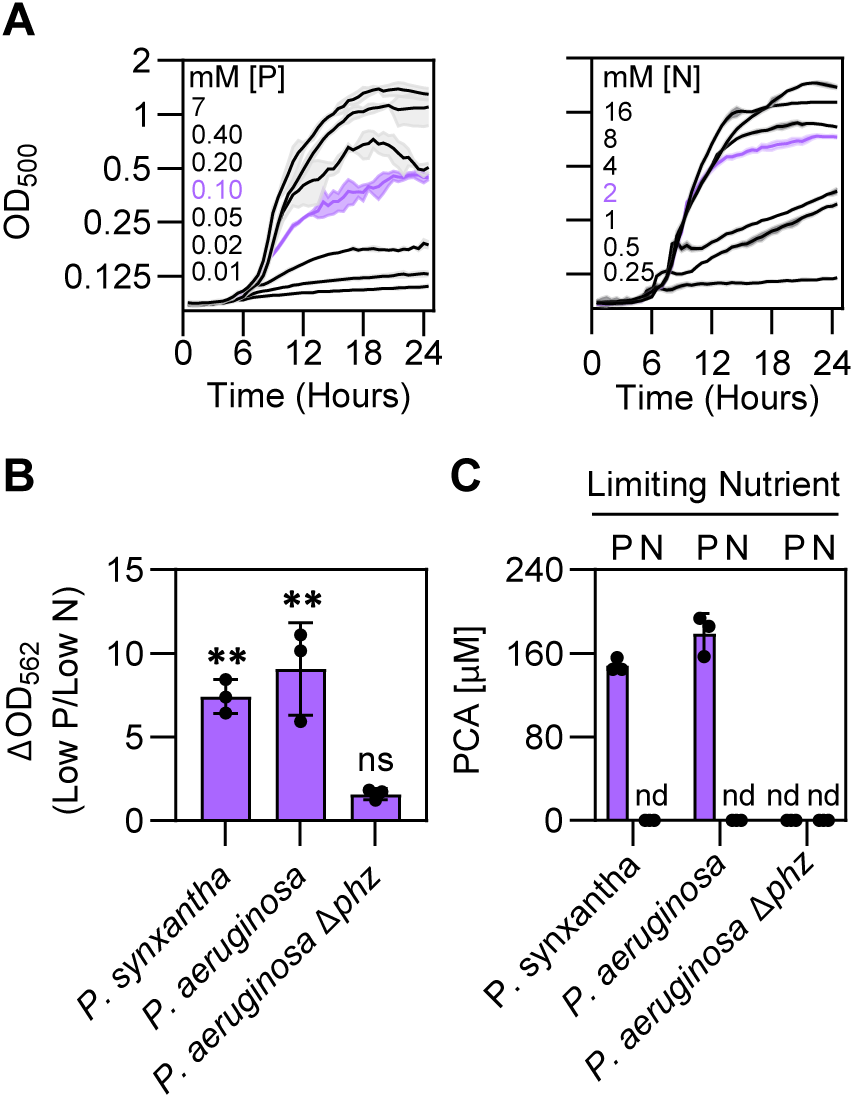
Development of the whole-cell phospho-ferrozine assay. **(A)** Effect of different phosphorus (left) and nitrogen (right) concentrations on *P. synxantha* growth. Phosphorus and nitrogen concentrations selected for the phospho-ferrozine assay are colored purple. **(B)** The ferrozine assay detects redox-active metabolite production in response to P limitation. Data shown are the ratios of the growth normalized ferrozine signal from P-limited strains (Low P) versus N-limited strains (Low N). See Figure S1 for ΔOD_562_ values for each condition. Statistical significance was determined by One-Way ANOVA analysis comparing ferrozine levels from the phosphorus-limited condition (low P) to the nitrogen-limited (low N) condition within each strain. ns = not significant; ** = p < 0.01. **(C)** Liquid-chromatography mass spectrometry data showing that the redox-active metabolite PCA is produced under P-limitation but not N-limitation. nd = not detected. **(A-C)** Data shown are the means ± SD of three biological replicates.

We next tested if the ferrozine assay could detect P-regulated RAM production from pseudomonads. Our assay depends on electron transfer from RAMs to iron(III) and therefore requires that RAMs be in their reduced, electron-carrying state (Fig. 1A). In the case of phenazines and pseudomonads, the mechanism of metabolite reduction is an open area of research but is thought to be facilitated by the bacterial cell (37, 38). In addition, phenazines are easily oxidized by oxygen in air, rendering them unable to perform iron(III) reduction (3, 39). To both facilitate phenazine reduction and minimize phenazine oxidation, we incubated cultures in an anaerobic chamber for one hour prior to performing the ferrozine assay. With this approach, the ferrozine signal in two phenazine-producing pseudomonads, *P. synxantha* and *Pseudomonas aeruginosa*, was significantly higher under P-limitation as compared to N-limitation (Fig. 2B and Fig. S1). Importantly, the ferrozine signal produced in cultures of a *P. aeruginosa* strain with deletions in phenazine biosynthetic genes (Δ*phz*, (40)), was low and did not change in response to P-limitation (Fig. 2B and Fig. S1). Using Liquid-Chromatography Mass-Spectrometry (LC-MS) we verified that our growth conditions led to changes in PCA production (Fig. 2C). P-limitation stimulated PCA production in both *P. synxantha* and *P. aeruginosa,* but not the *P. aeruginosa* Δ*phz* strain. No PCA was detected when any strain was limited for N (Fig. 2C). Altogether, these experiments indicate that the ferrozine assay in combination with P-limitation can reliably be used to screen for bacteria that produce RAMs.

### Application of the phospho-ferrozine assay to a diverse panel of root-associated bacteria

To extend our findings beyond strains in current culture collections, we generated a library of root-associated bacteria. Isolates were obtained from three sites and 20 plants collected across the United States: *Brachyelytrum aristosum* grasses in Harvard Forest, *Panicum virgatum* grasses in Konza Prairie Biological Station, *Digitaria californica* grasses in Santa Rita Experimental Range, and commercial tomato plants (Fig. 3A and B, Table S1 and Table S2). In preliminary experiments using commercial tomato plants, we used 16S amplicon sequencing to compare the effects of isolation media supplemented with either single carbon sources (glucose, succinate, or pyruvate), or yeast extract on the diversity of bacterial isolates. As expected, diversity decreased between whole soil and plates across media types. However, yeast extract maintained the highest level of diversity (Fig. S2) and was used for all further experiments (Fig. 3C and Fig. S3). A total of 557 strains of at least 48 genera were collected across all sampling sites. As in preliminary tests, our isolation method primarily enriched for Proteobacteria, Actinobacteria, Firmicutes, and Bacteroidetes (Fig. 3C, Fig. S3, and Table S1).

**Figure 3.**
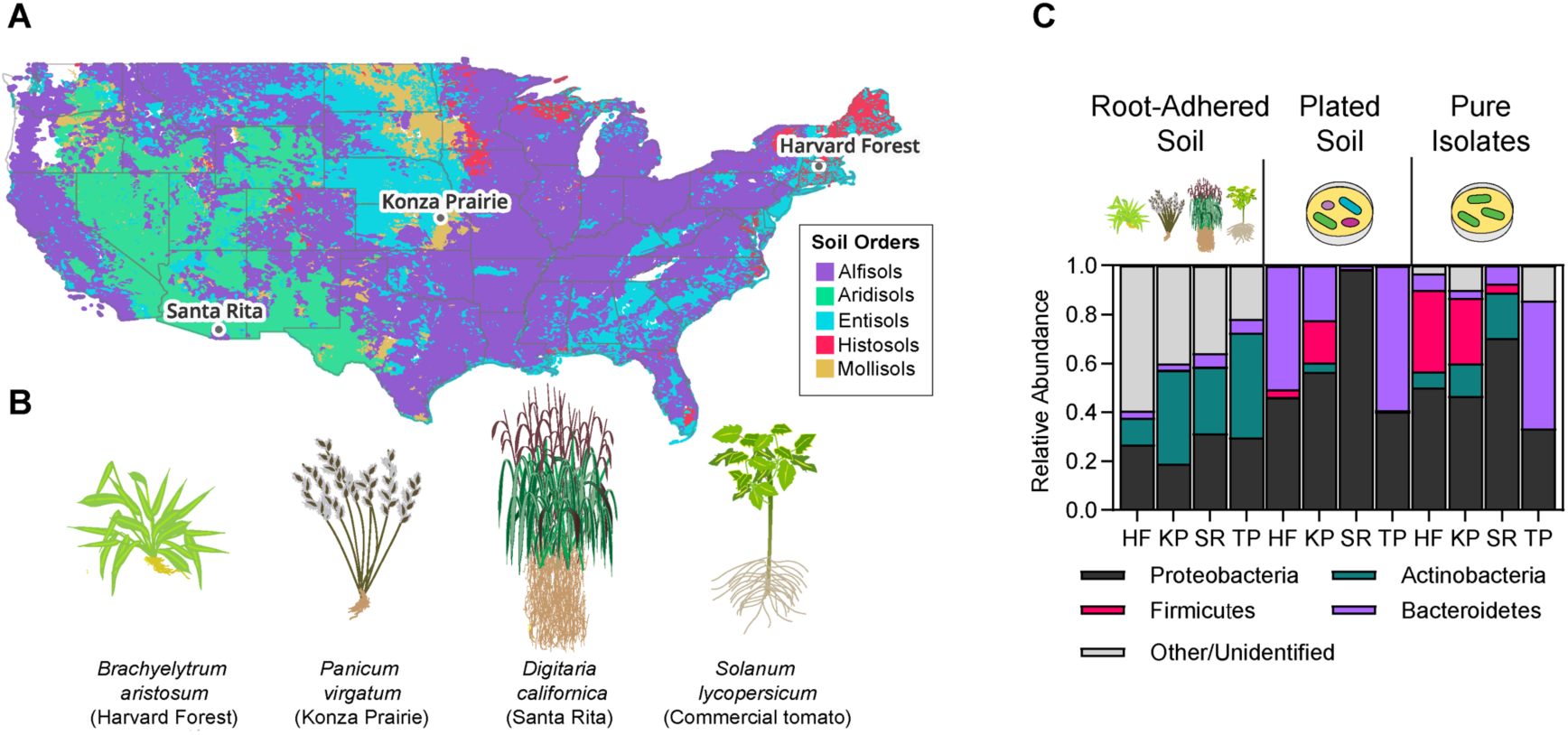
Isolation of root-associated bacteria from multiple geographic locations. **(A)** Sites where plants were collected. Map is colored by soil order and was generated using QGIS. See Table S2 for GPS coordinates and harvest dates. **(B)** Plant species collected. Five individual plants growing in close proximity were collected at each location. **(C)** 16S rRNA amplicon sequencing of bacteria from root-adhered soil (left), yeast extract plates (middle), and isolated strains (right). The data shown represent one plant per site. See Figure S4 for data from replicates. HF = Harvard Forest; KP = Konza Prairie; SR = Santa Rita; TP = commercial tomato plants.

We used the phospho-ferrozine assay to screen all 557 soil isolates for the ability to produce RAMs in response to P-limitation. 101 soil isolates showed poor growth (OD_562_ < 0.10); and were excluded from further analysis. P-limitation increased the ferrozine signal by at least 2.0-fold in 128 of the remaining 456 (28%) soil isolates (31 Harvard Forest isolates, 34 Konza Prairie isolates, 33 Santa Rita isolates, and 30 tomato plant isolates, Fig. 4A and Table S3). At least 19 genera across Proteobacteria, Actinobacteria, Firmicutes, and Bacteroidetes showed an increased ferrozine signal in response to P-limitation (Fig. 4B and Table S3B). The majority of the positive hits identified were Proteobacteria (61%), followed by Actinobacteria (11%), Bacteroidetes (6%), and lastly, Firmicutes (5%) (Fig. 4B and Table S3B). Despite multiple attempts to PCR amplify the 16S region, 21 positive hits remained unidentified (Fig. 4B and Table S3). The highest phospho-ferrozine signal for each location were from isolates identified at the genus level as *Citrobacter* (tomato plants) *Stenotrophomonas* (Harvard Forest), *Pseudomonas* (Konza Prairie) and *Bacillus* (Santa Rita). Isolates from Konza Prairie and Santa Rita showed the highest overall phospho-ferrozine signal, increasing by ∼20 fold when limited for phosphorus (Fig. 4A). These results suggest that a diverse array of soil bacteria across multiple phyla can produce RAMs in response to P-limitation.

**Figure 4.**
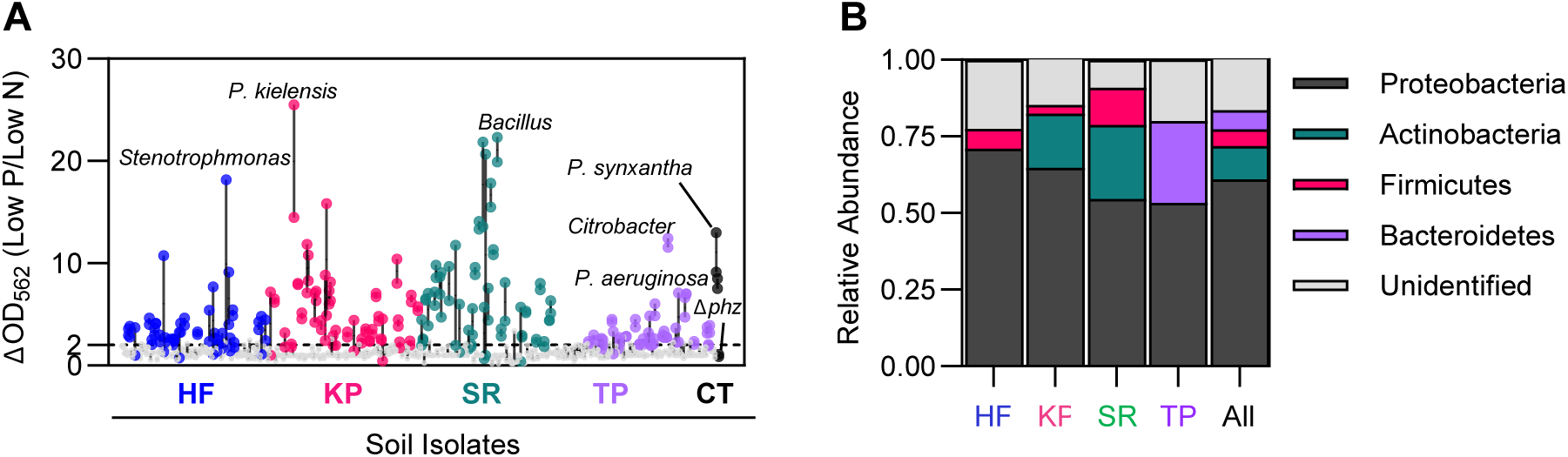
A diverse panel of soil isolates produce redox-active metabolites in response to phosphorus limitation. **(A)** Identification of RAM-producing bacteria with the phospho-ferrozine screen. 557 isolates were screened, and 101 soil isolates that did not grow (OD562 < 0.10) were excluded. Two replicates of the remaining 456 soil isolates are shown, comparing the ferrozine signal from P-limited (Low P) to N-limited conditions (Low N). Replicates for each strain are connected by a line. Soil isolates with an average ΔOD_562_ ratio ≥ 2.0 are considered positive hits and are colored by the site they were isolated from. Soil isolates with an average ΔOD_562_ ratio < 2.0 are depicted in grey. See Table S3A for ferrozine signal data of all the tested soil isolates. **(B)** Taxonomic distribution of the 128 soil isolates with an average ΔOD_562_ ratio ≥ 2.0. Genera were assigned based on a BLAST search of the amplified 16S rRNA sequence and soil isolates were classified by phyla. See Table S3B for a list of the identified positive hits from the phospho-ferrozine screen. **(A and B)** Soil isolates are organized by site. HF = Harvard Forest; KP = Konza Prairie; SR = Santa Rita; TP = commercial tomato plants; CT = controls.

### Verification of diffusible redox-active metabolite production using cyclic voltammetry

When conducted with whole cells, the phospho-ferrozine screen cannot distinguish between iron(III) reduction by cellular machinery or cell-associated metabolites vs iron(III) reduction by diffusible RAMs. As our primary interest was in the latter, we investigated whether the ferrozine signal we observed from whole cells was in fact due to diffusible metabolites by conducting the phospho-ferrozine assay on filter-sterilized supernatants from a subset of P-limited soil isolates. We prioritized isolates that showed a strong response in the whole-cell assay and aimed to capture broad taxonomic diversity. Of the 48 soil isolate supernatants tested, the ferrozine signal was significantly increased under low P in 22 soil isolate supernatants (Fig. 5A and Table S4). Supernatants from a Konza Prairie *Pseudomonas* isolate showed the highest ferrozine signal, which was 8.1-fold greater than *P. aeruginosa* Δ*phz*. (Fig. 5A). These results suggest that the whole-cell ferrozine assay detects iron(III) reduction by diffusible RAMs in our soil isolates.

**Figure 5.**
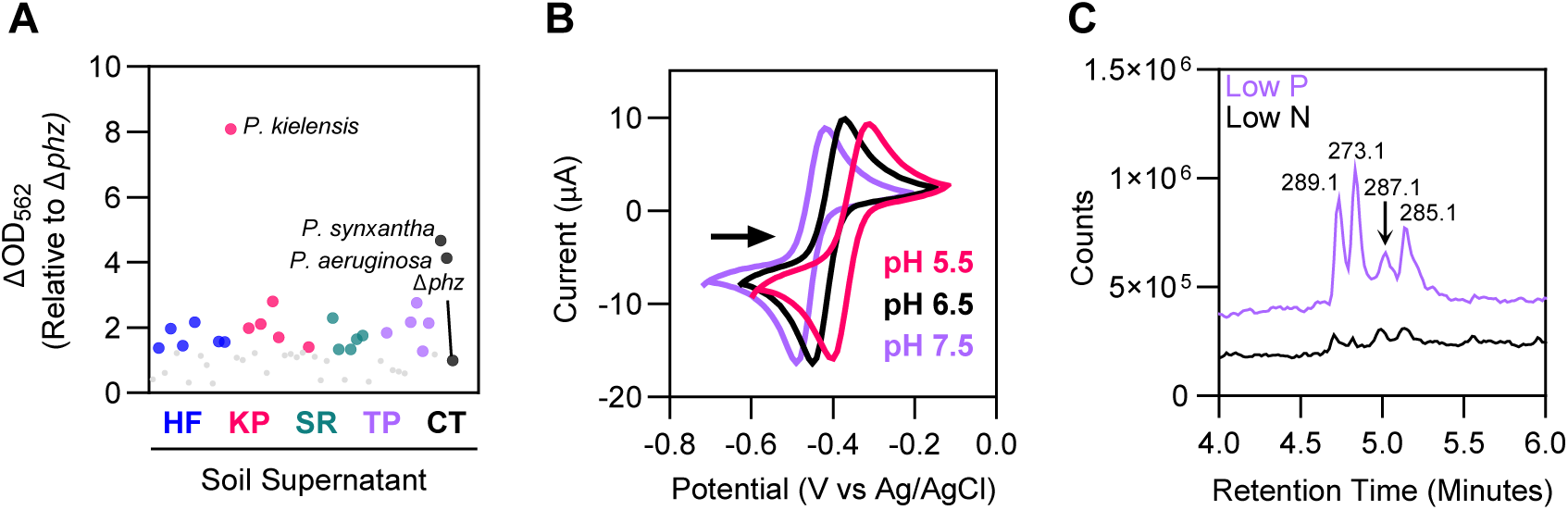
The phospho-ferrozine assay detects diffusible redox-active metabolites. **(A)** Redox-active metabolites are secreted extracellularly in a subset of soil isolates. Phospho-ferrozine assay of filter-sterilized supernatants from P-limited soil isolates. Data shown are the means of two replicates comparing the ferrozine signal of each strain to *P. aeruginosa* Δ*phz*. Supernatants with statistically significant higher ferrozine signal compared to *P. aeruginosa* Δ*phz* are colored by site (p < 0.05). No significant increase = grey (p > 0.05). Statistical significance was determined with a One-way ANOVA test. HF = Harvard Forest; KP = Konza Prairie; SR = Santa Rita; TP = commercial tomato plants; CT = controls. See Table S4 for supernatant ferrozine data. **(B)** Voltammogram showing the presence of a small molecule with reversible, proton-coupled redox activity in P-limited *P. kielensis* supernatant at pH 5.5, 6.5, and 7.5. Redox potentials: pH 5.5 = -153 mV; pH 6.5 = -203 mV; pH 7.5 = -248 mV (vs NHE). Scan shown is the second of three scans and a gold working electrode was used. **(C)** Total ion chromatogram of *P. kielensis* supernatants showing several putative small molecule masses produced under phosphorus but not nitrogen limitation.

To further confirm the presence of redox-active molecules in supernatants, we employed cyclic voltammetry, a standard electrochemical method often used to characterize redox activity (41). Using cyclic voltammetry, we were clearly able to detect the presence of phenazines in supernatants from P-limited wild type *P. synxantha* and *P. aeruginosa* cultures, but we did not observe any signal in *P. aeruginosa* Δ*phz* strains (Fig. S4). We also detected redox-active compounds in the supernatants of multiple soil isolates including *Chromobacterium*, *Paenarthrobacter, Arthrobacter* and *Pseudomonas* (Fig. S4). However, not all supernatants that showed an elevated phospho-ferrozine signal showed clear evidence for the presence of redox-active compounds (Fig. S4), potentially due to concentrations being too low or molecule damaged during drying. Of note, a *Pseudomonas* isolate from Konza Prairie showed robust P-regulated RAM production from whole-cell and supernatant ferrozine assays as well as cyclic voltammetry analysis (Fig. 5B and Fig. 4S); we chose to focus further efforts on understanding RAM production in this organism.

### *Pseudomonas kielensis* produces a non-canonical redox-active metabolite under low phosphorus

To investigate the biosynthetic capacity of our *Pseudomonas* isolate, we conducted Illumina whole genome sequencing. Using a full length 16S sequence, this isolate was identified to the species level as *Pseudomonas kielensis* with 99.4% 16S identity to its closest relative by best BLAST. As pseudomonads frequently produce phenazines, we suspected this organism might be synthesizing some variant of this type of RAM. However, a search of the genome using antiSMASH (42) failed to identify any phenazine biosynthetic gene clusters. A BLAST search using the phenazine biosynthetic genes *phzA-F* from *Pseudomonas fluorescens* (tblastn, e-value cutoff of 0.001) revealed only one hit to a gene annotated as *phzF*, which was not co-located with other phenazine genes (Fig. S6A) suggesting that this organism does not encode phenazine biosynthetic gene clusters. To check the functionality of the *phzF* homolog, we created a clean deletion of the gene. The Δ*phzF* mutant showed a small decrease in the phospho-ferrozine signal but the majority of the signal was maintained (Fig. S6B), suggesting that this gene was not a major contributor to RAM production.

To more directly investigate possible phenazine production, we determined the redox potential of the putative metabolite and used untargeted metabolomics to analyze supernatants from cultures grown under high and low P. Cyclic voltammetry experiments conducted across pH values (5.5-7.5) showed that the oxidation and reduction peaks from the *P. kielensis* supernatant shifted with pH, consistent with a proton-coupled redox reaction (Fig. 5B). However, the calculated midpoint potential of the *P. kielensis* supernatants ranged from -153 mV to -248 mV (vs NHE), which is far more negative than the potentials reported for PCA across this pH range (-1mV to -113 mV vs NHE) as well as most well-studied phenazines (3, 39). We used metabolomics to search for phenazines and other metabolites in N- and P-limited *P. kielensis* supernatants (see Fig. S5 for characteristic growth). We were unable to detect any phenazines using searches for the masses of PCA, pyocyanin, 1-hydroxy-phenazine, or phenazine-1-carboxamide (m/z = 225.2, 211.2, 197.2, 224.2, ([M+H]^+^). A comparison of *P. kielensis* supernatants to purified standards of the phenazines pyocyanin and PCA also showed no clear overlap. Instead, our untargeted analysis revealed several peaks in the total ion chromatogram that were present under low P but not low N (Fig. 5C). These corresponded to m/z =273.1, 285.1, 287.1 and 289.1 ([M+H]^+^). All four peaks were also present in supernatants from the Δ*phzF* mutant (Fig. S7), further supporting the hypothesis that this gene does not contribute to RAM production. Altogether, our data suggest that *P. kielensis* produces a RAM with reversible and proton-coupled redox activity, but that this molecule is distinct from well-known phenazines, possibly representing a different class of compounds. Investigations into the structures and biosynthetic pathways of these molecules are currently underway.

### Application and broader implications of the phospho-ferrozine assay

Here we show that the phospho-ferrozine assay detects the presence of RAMs in pseudomonads known to make these molecules and that this screen can be applied to a variety of soil isolates. As in any screen, the fidelity between the observed signal and RAM production must be independently verified. One key concern is the possibility of false positives resulting from cellular iron reduction instead of RAM production. Our experiments with *P. aeruginosa* phenazine-null mutants show that in this organism, non-specific cellular iron(III) reduction does not lead to a positive ferrozine signal (Fig. 2B). Nonetheless, it is notable that in some cases cell-free supernatants showed a drop in ferrozine activity or failed to show a clear signal when analyzed with cyclic voltammetry. As we only performed cyclic voltammetry on samples that showed positive supernatant ferrozine assays, the lack of small molecule detection via this method is most likely due to low concentrations of RAMs or molecule damage during sample dry-down and resuspension. The discrepancy between whole-cell and supernatant ferrozine assay is more complex. One potential culprit could be dissimilatory iron reduction, a specialized anaerobic metabolism in which iron(III) replaces oxygen as the terminal electron acceptor (43). However, this process is not known to be regulated by P and is typically repressed by oxygen (44). Our experiments were conducted under aerobic conditions with only a one-hour anaerobic incubation in which isolates might activate dissimilatory iron reduction machinery, making it unlikely that this process is a major contributor to false positives. Another consideration is that in order to reduce iron(III), RAMs must be maintained in their reduced state and this process may require cellular reduction (37, 38). As such, instead of false positives in our whole-cell assays, some of our supernatant results may be false negatives driven by incomplete metabolite reduction. It is also possible that some of our isolates produce RAMs that are tightly cell-associated and are removed during filtration. Our primary interest has been in diffusible metabolites, but the screen could easily be adapted to detect these types of cell-associated RAMs.

While care must be taken to validate results and some false positives and negatives are to be expected, the phospho-ferrozine screen represents a rapid, high-throughput, and relatively inexpensive way to screen for RAM production. The screen is comparable to the classic Chrome-Azurol-S (CAS) Assay (45) which selects for a specific chemical feature (iron binding) and uses nutrient (iron) stress to stimulate production. Independent verification is also needed for the CAS Assay, but it has been of extraordinary utility in detecting siderophores and in activity-guided studies to isolate these molecules. We expect the phospho-ferrozine screen to facilitate similar advances in isolation and characterization of RAMs as well as studies to determine the prevalence and patterning of P-regulated RAM production across microbial communities. A link between secondary metabolites and P stress has been well documented in the literature (4, 25). However, previous studies have mostly focused on strains in culture collections, leaving open the question of whether this is truly widespread in soils. Our finding that 28% of isolates produced a positive phospho-ferrozine signal suggests that this feature is common across genera and environments. Small modifications to the screen such as storing isolate libraries in an array format in 96-well plates could allow the assay to be applied at a very large scale and open the door for more biogeographic studies of probing the distribution and frequency of RAM production across environments. Indeed, previous studies focused on phenazines have shown that the capacity to synthesize these molecules can be present in 1-2% of bacteria (36, 46). Future studies that expand the search to all RAMs and incorporate regulatory logic and soil phosphorus availability will be an exciting next step for the study of RAMs as well as secondary metabolites in general.

Finally, our findings underscore the importance of considering nutrient stress when studying secondary metabolites and highlight an opportunity to fuse these types of studies with investigations of small molecule elicitors of secondary metabolite biosynthesis. As shown here and elsewhere, phenazine biosynthesis in pseudomonads is frequently regulated by P stress. However, as is common for secondary metabolites, phenazines are also regulated by small molecule elicitors (47) including quorum sensing molecules. This type of combined nutrient and small-molecule-based regulation occurs for siderophores (48–50) and we expect it will also occur in our newly discovered RAM producers. In the future, it will be important to blend studies of small molecule and nutrient stress regulation. While challenging, this endeavor holds great promise for understanding the environment experienced by microbes in the wild and will contribute to applied efforts to manipulate microbes across biomedical and soil contexts. Such combinatorial studies may aid the discovery of novel metabolites that might act as therapeutics and help us understand when and how they might be produced. There is also growing momentum for using secondary metabolites to manage microbial functions in ways that promote plant health – either through the suppression of pathogens or enhancement of nutrient access (51–53). Phosphorus bioavailability is of paramount importance in natural and agricultural systems (27, 54, 55) and our findings that numerous bacteria produce RAMs in response to P stress bodes well for future work aimed at understanding P cycling in natural systems and leveraging these small molecules in managed ones.

## Conclusion

We developed a tool to rapidly identify RAM-producing bacteria from the soil by utilizing P limitation to induce RAM biosynthesis and measuring redox activity through the ferrozine assay. We show that multiple genera across four different phyla secrete RAMs in response to P limitation, suggesting this regulatory mechanism is widespread. Further, we show that despite the frequency of phenazine biosynthesis in pseudomonads, *P. kielensis* most likely does not synthesize these metabolites and instead makes a different RAM. Our work opens the door for studies investigating how the production of these small molecules is distributed across different environments, the multi-layered molecular mechanisms microbes use to regulate their biosynthesis, and the overall role RAMs play within microbial communities living at plant roots.

## Materials and Methods

### Bacterial strains and growth conditions

Strains, plasmids, and primers used in this study are listed in Table S5. A list of soil isolates and their closest 16S identity is provided in Table S1. Unless stated otherwise, all bacterial strains were grown at 30 °C on Luria Broth (LB) agar plates prior to experiments. Liquid cultures were grown at 30 °C with continuous shaking at 200 RPM except for *Escherichia coli* strains which were grown at 37 °C.

Bacterial strains were grown in defined media, (8.22 mM glucose, 12.33 mM succinate, 16.44 mM pyruvate, 16 mM NH_4_Cl, 4.70 mM K_2_HPO_4_, 2.30 mM KH_2_PO_4_, 0.68 mM CaCl_2_, 0.41 mM MgSO_4_, and Aquil trace metals (56) supplemented with 10 µM iron, buffered at pH 5.5 with 25 mM MES). For P limitation, P was adjusted to 100 µM (K_2_HPO_4_ = 33 μM and KH_2_PO_4_ = 67 μM) while N was maintained at 16 mM. For N limitation, NH_4_Cl was adjusted to 2 mM and P was maintained at 7 mM. For growth gradient experiments, cultures were grown overnight in defined media, washed and resuspended at OD 1.0 (500 nm) in either P-free or N-free defined media, and inoculated at an OD_500_ of 0.01 in 96-well plates. Growth was tracked via absorbance at 500 nm every 30 minutes for 24 hours using a BioTek Epoch 2 microplate reader with constant shaking.

### Abiotic ferrozine assays

Abiotic ferrozine assays were performed in potassium chloride (KCl) buffer (1 M KCl buffered with 10 mM ammonium acetate-MOPS at pH 6) using a final concentration of 200 μM FeCl_3_ and 2 mM ferrozine and the ferrozine-Fe(II) complex was quantified using absorbance at 562 nm.PCA was reduced electrochemically under an N_2_ headspace as previously described (3) and experiments were performed anaerobically (95% N_2_, 5% H_2_) to minimize PCA oxidation by oxygen.

### Liquid-chromatography mass-spectrometry

For metabolomics studies, filtered supernatants were analyzed directly on an Agilent 6125B single quadrupole mass spectrometer fitted with a diode array detector. Briefly: 2 µl of sample was injected into a C18 column (Agilent Poroshell 120: 50mm length, 2.1 mm internal diameter, 2.7 µm particle size). Separation was achieved using a gradient from acidified (0.1% formic acid) 90% water/10% acetonitrile to 100% acetonitrile over 6 minutes. PCA and pyocyanin peaks were identified by comparison to purchased standards (Sigma) and quantified using a standard curve and the extracted absorbance at 315 nm.

### Isolation of root-associated bacteria from plants

Five individual plants of the grass species *Brachyelytrum aristosum*, *Panicum virgatum*, and *Digitaria californica* were collected from National Ecological Observatory Network (NEON) sites at Harvard Forest, Konza Prairie, and Santa Rita, respectively (see Table S2 for GPS coordinates and sampling). “Big Beef” variety tomato plants were purchased. NEON samples were transported on ice and processed within three days of harvest. To isolate root-associated bacteria, soil was shaken off the plant until only soil adhered to plant roots remained. Roots were cut near the base of the stem, placed in conical tubes with sterile wash buffer (0.40 mM MgSO_4_ and 1.7 mM NaCl, pH 6.0), vortexed for 10 seconds, sonicated for one minute, and vortexed again for an additional 10 seconds. The resulting slurry was diluted 10-100X and plated onto agar plates. Preliminary experiments used plates that contained glucose, succinate, or pyruvate as sole carbon sources (10 mM) or 1 g/L yeast extract and 1g/L tryptone in addition to 0.5 mM NH_4_Cl, 0.5 mM K_2_HPO_4_, 1 mM MgSO_4_, 2 mM NaCl, MEM essential amino acids (1.6 ml/L, Sigma M5550), Aquil trace metals (56) supplemented with 10 µM Fe, and 50 μg/mL nystatin and were incubated at 25°C for 48 hours. Yeast extract plates were used for all subsequent analyses. Colonies that appeared morphologically distinct were picked and re-streaked three times to purity before being freezer stocked. To ensure the sterility of the process, sterile wash buffer was also plated and showed no microbial growth.

### 16S amplicon sequencing of soil and plate samples

Bacterial diversity on plates was determined by scraping the entire plate. Plates were incubated for 24 hours, 2 ml phosphate-buffered saline solution (8 g/L NaCl, 0.2 g/L, 1.44 g/L Na_2_HPO_4_, 0.24 g/L KH_2_PO_4_) was applied to the plate, cells were homogenized using a sterile scraper, collected in an Eppendorf tube, centrifuged at 10,000xg for five minutes and pellets were stored at -80°C. For soils, a slurry (described above) was centrifuged at 10,000xg for 20 minutes, supernatants were decanted, and soil pellets were stored at -80°C.

Genomic DNA from plates and soil samples was extracted using the DNeasy Blood & Tissue Kit (Qiagen) and the DNeasy PowerLyzer Power Soil Kit (Qiagen), respectively. 16S paired-end amplicon sequencing was performed by SeqCenter (Pittsburgh, PA, USA) using the 341F (CCTACGGGDGGCWGCAG, CCTAYGGGGYGCWGCAG) and 806R(GACTACNVGGGTMTCTAATCC) primers. Amplicons were sequenced on a P1 600cyc NextSeq2000 Flowcell. Bioinformatic analysis was conducted using a standard QIIME2 pipeline (57). Adapters were removed using CutAdapt. Denoising and trimming was performed with DADA2: forward and reverse reads were trimmed to 250 bp, >100,000 reads were analyzed for each sample, except for tomato plant plates and soils samples where only ∼60-77K reads were retained. Taxonomy was assigned using the Silva classifier. 16S amplicon sequences were deposited in Sequence Read Archive under project number PRJNA1152629.

### Identification of soil isolates

The taxonomic identity of purified isolates was determined via colony PCR. Colonies resuspended in nuclease-free water and lysed by sonicating for five minutes at 40 kHz, followed by boiling for 10 minutes at 100 °C. Cell lysates were pelleted and clarified lysates were used for PCR reactions. The 16S rRNA gene was amplified using 1492R (TACGGYTACCTTGTTACGACTT) and 27F (AGAGTTTGATCMTGGCTCAG) primers (58). PCR products were run on an agarose gel to confirm the presence of a single amplicon at the expected size (∼1,500 bp) and Sanger sequenced with the 27F primer at Epoch Life Science (Missouri City, TX, USA). Strain identity was obtained through a BLAST search against the NCBI 16S rRNA sequence database (Table S1). 220 of 370 16S amplicon sequences were deposited in GenBank under the accession numbers PQ223443-PQ223663. The remaining 16S amplicon sequences failed to meet the requirements for submission to GenBank.

### Whole-cell phospho-ferrozine screen

*Pseudomonas* cultures or soil isolates were grown for 24 hours in 200 μl P- and N-replete media in a 96-well plate, transferred (1:100) to new 96-well plates containing either 200 μl P-limited or N-limited media, and grown for an additional 24 hours. Plates were then incubated statically for one hour in an anaerobic chamber (95% N_2_ and 5% H_2_) to remove residual oxygen (3, 39). Subsequently, 500 μM of FeCl_3_ (maintained as a concentrated stock in 0.1 N HCl) followed by 2 mM ferrozine (in 2 M MOPS, pH 7) was added, and the ferrozine-Fe(II) complex was detected immediately via absorbance at 562 nm using a BioTek Synergy HTX plate reader. Ferrozine stock solutions were degassed in the anaerobic chamber for at least three days prior to experiments. To account for differences in growth that might contribute to the OD_562_ signal, absorbances at 562 nm for each isolate were recorded prior to the addition of FeCl_3_.This background number was subtracted from the final ferrozine signal (denoted as ΔOD_562_). The ferrozine signal for each soil isolate is provided in Table S3.

### Supernatant phospho-ferrozine assays

Cultures were grown overnight in defined media, diluted to an OD_500_ of 0.01 in 10 ml P-limited media and grown for and additional 24 hours. P-limited cultures were then incubated statically for one hour in an anaerobic chamber to deplete oxygen, and supernatants were filter sterilized using a 0.22 μm Spin-X column (Corning). 200 μl of filter-sterilized supernatant was transferred to a 96-well plate and mixed with 500 μM FeCl_3_ and 2 mM ferrozine. After 40 minutes, the ferrozine-Fe(II) complex was detected at 562 nm (BioTek Synergy HTX plate reader). The remaining volume of supernatant was used for cyclic voltammetry experiments below.

### Cyclic voltammetry

Supernatants from P-limited cultures were filter-sterilized (0.22 μm), concentrated under a stream of N_2_ gas, and resuspended in 1 ml KCl buffer. Cyclic voltammetry measurements were conducted using a BioLogic SP-300 potentiostat at a scan rate of 50 mV/s using a platinum wire counter electrode, an Ag/AgCl reference electrode and a 3 mm gold working electrode. All experiments with coumarin and PCA were conducted with 1mM solutions in KCl buffer. pH gradient experiments were conducted under a constant stream of N_2_ gas. Midpoint redox potentials were calculated as the average of the potentials at minimal and maximal current (vs Ag/AgCl) + 207 mV for comparison to the NHE.

### Genomic sequencing and annotation

*P. kielensis* genomic DNA from an LB liquid overnight culture was extracted using the DNeasy Blood & Tissue Kit (Qiagen). Sequencing, assembly, and gene annotation were performed by SeqCenter (Pittsburgh, PA, USA) using an Illumina NovaSeq X Plus sequencer. The *P. kielensis* genomic sequence was deposited in GenBank under the accession JBGRQV000000000. The version described in this study is version JBGRQV010000000.

### Construction of *P. kielensis* deletion strains

In-frame deletions were constructed by allelic exchange using the pMQ30 shuttle vector containing ∼1 kb flanking regions which were built by Gibson Assembly (59). Colony PCR and Sanger sequencing were used to confirm plasmid construction. Plasmids were introduced to *P. kielensis* by triparental conjugation and merodiploids were selected on Vogel-Bonner (VBMM) plates containing 50 μg/mL gentamycin (60, 61). Transconjugants were counter selected on 5% sucrose LB medium with no added salt. Deletions were confirmed by colony PCR.

## Acknowledgments

We thank members of the McRose lab for insightful and helpful discussions and R. Shafiee for help with QGIS. This work was supported by the James Foundation, startup funds from the MIT Department of Civil and Environmental Engineering and a L’Oréal USA for Women in Science Fellowship granted to DLM. Sample collection was conducted in collaboration with the National Science Foundation’s National Ecological Observatory Network (NEON) program.

